# Distinct lipid transport proteins are regulated by innate immune stimuli

**DOI:** 10.64898/2026.01.11.698848

**Authors:** Lydia P. Tsamouri, Najd M. Aljadeed, Antoni Olona, Paras K. Anand

## Abstract

Lipid transport plays a critical role in the distribution of lipids across subcellular compartments. This is pivotal during infection and other stress stimuli that increase metabolic demands. While lipid biosynthesis is regulated by immune stimuli, whether immune signalling also influences lipid transport mechanisms remains unexplored. Here, we demonstrate that TLR signalling impacts the gene expression of lipid transport proteins in human monocytic THP-1 cell line and compare it to the expression in primary bone marrow-derived mouse macrophages. Our data demonstrates that TLR4 signalling selectively modulates the expression of oxysterol-binding protein-related proteins (ORPs), a key family of proteins that transport lipids between organelles. Remarkably, TLR4 activation led to the downregulation of several ORP family members in human THP1-derived macrophages. However, this response was less profound in mouse macrophages. In contrast, the expression of steroidogenic acute regulatory domain (STARD) proteins, many of which transport lipids between mitochondria and other compartments, remained broadly unaffected. Moreover, IFNG, a cytokine that plays a key role in the immune response, did not considerably impact human ORP or STARD expression levels, either alone or in combination with LPS. Together, these results reveal that TLR signalling exerts selective and critical control over lipid trafficking pathways with important species-specific differences. These findings provide new insights into the crosstalk between immune signalling and lipid metabolism, which may offer novel targets to treat diseases characterized by dysregulated lipid pathways.

## Introduction

Lipids play a fundamental role in diverse cellular functions supporting the structural and energy needs of all cells [1,2]. Additionally, lipids are critical to intracellular signalling and serve as second messengers. In particular, lipid metabolism in immune cells is dynamically rewired in response to cues sensed during infection and other stress stimuli [1,3]. For example, cytokines like IFNG, a key modulator in immune responses, have been shown to influence lipid metabolism by decreasing cholesterol efflux and modulating lipid availability, thereby shaping immune cell activation and function [4,5]. These shifts in metabolic pathway allow immune cells to meet the increased demands for effective defence[6]. They also help counter homeostatic imbalances during infection and stress. Consequently, lipids not only support cellular signalling but also reprogram in response to immune cues to enhance host defence, drive inflammatory signalling, and maintain homeostasis.

While the impact of TLR signalling on lipid metabolism has been explored, it remains unclear whether immune signalling directly influences the mechanisms governing lipid trafficking, particularly through lipid transport proteins. Given their essential roles in cellular function, lipid transport mechanisms are integral to immune cell activity. These pathways maintain lipid distribution across compartments, enabling quick metabolic adjustments when required. Moreover, lipid metabolism is being increasingly recognized as a key target influenced by immune signalling, including through the TLR pathway [7]. TLR recognition of pathogen-associated molecular patterns initiates cascades leading to the activation of NFKB and MAP-kinase transcription factors, which in turn regulate the expression of a variety of genes [8]. Published research indicates that TLR signalling influences lipid metabolism in varied ways [9–11]. TLR ligation modulates pathways that activate SREBPs and LXRs, key transcription factors involved in lipid metabolism [12–15]. These transcription factors govern lipid biosynthesis and efflux, orchestrating the size of cholesterol pool based on cellular demand. This is particularly evidenced is case of TLR4 ligation by bacterial LPS resulting in increased cholesterol biosynthesis to support the high lipid demands of immune cells under stress [16,17]. Thus, there is a significant association between lipid metabolism and immune pathways. This crosstalk underlies the pathogenesis of several infectious and metabolic diseases. However, it remains unclear if immune signalling has a direct impact on the mechanisms that facilitate lipid movement between cellular compartments.

Here, we wanted to investigate whether TLR signalling differentially regulates lipid transport proteins, focusing on the oxysterol-binding protein-related proteins (ORPs) and steroidogenic acute regulatory domain (STARD) proteins. Understanding these mechanisms may reveal novel therapeutic targets for immune and metabolic diseases. The transfer of lipids across membranes is predominantly dependent on vesicular transport, lipoprotein-mediated transport, and lipid transfer proteins [18,19]. Lipid transport proteins, in particular, are essential for the movement of lipids at membrane contact sites where organelles come close together but may not physically fuse. The major lipid transport protein families include oxysterol-binding protein-related proteins (ORPs), steroidogenic acute regulatory domain (STARD) proteins, and GRAMD proteins [18]. The ORP proteins are a key family of 12 members that mediate the non-vesicular transport of sterols and phospholipids between organelles, such as the ER and the plasma membrane [20][19,20]. ORPs are classified into six subfamilies (I-VI) based on structural similarities and domain architecture (**Fig. S1**). Each ORP features a conserved oxysterol-binding protein-related domain (ORD), responsible for their lipid transport function, and additional accessory domains that mediate localization and specificity. For example, pleckstrin homology (PH) domains enable ORPs to tether to specific lipid pools within membranes by recognizing phosphoinositide lipids. Several ORPs contain FFAT (two phenylalanines in an acidic tract) motifs, which facilitate interactions with the ER membrane. Additionally, ORP5 and ORP8, contain transmembrane (TM) domains that anchor them to specific organelles, enhancing their functional versatility. The diversity in lipid-binding preferences and the presence of accessory motifs confers unique roles in cellular lipid homeostasis [21,22]. The STARD protein family, in contrast, primarily facilitates intracellular cholesterol movement to mitochondria, supporting synthesis and energy metabolism [23,24]. The concerted effort of lipid transport proteins ensures precise distribution of lipids across cellular compartments, enabling cellular adaptability to metabolic demands and external stimuli.

Given the central role of lipid transport proteins in maintaining cellular lipid homeostasis, understanding how TLR signalling modulates these proteins is critical to uncovering the broader impact of immune signalling on lipid metabolism and its implications for disease. There is evidence suggesting that the interaction of TLR signalling with lipid transport is relevant to immune cell function [25]. For example, ORP proteins have been shown to modulate key proteins involved in cholesterol efflux from cells [26]. Despite these preliminary suggestions, whether TLR signalling regulates lipid transfer proteins remains unexplored. In this study, we demonstrate that TLR signalling specifically regulates the gene expression of ORP proteins. By contrast, the regulation of STARD family members remains unaffected. Furthermore, we identify species-specific differences in lipid transport regulation. Our findings expand our understanding of immune-regulated lipid metabolism which has the potential to identify therapeutic targets for lipid-related disorders and metabolic diseases.

## Materials and methods

### Ethical statement

All procedures on animals were performed under appropriate personal and project licenses (PPL number: PP4111881). The procedures were carried out in accordance with the Animals (Scientific Procedures) Act 1986 and were approved by the Imperial College Animal Welfare and Ethical Review Body (AWERB) and the UK Home Office.

### Macrophage cell culture

WT C57BL/6J mice were obtained from Charles River, UK. The *in vitro* experiments used male and female mice in similar proportions, ranging in age from 8 to 12 weeks old. Bone marrow cells were obtained from the femurs and tibias from the mice and allowed to differentiate into bone marrow-derived macrophages as described previously [27,28]. THP-1 monocytic cell line (ATCC, Catalog #: TIB-202; RRID: CVCL_0006) was maintained in RPMI-1640 medium containing 10% FBS, 2 mM L-glutamine, 1% penicillin-streptomycin, 4.5 g/L glucose, 1% HEPES, and 0.1% 2-mercaptoethanol. THP-1 cells were differentiated into macrophages by culturing them in RPMI-1640 medium supplemented with 20 nM phorbol-12-myristate-13-acetate (PMA) for 24 h.

Peripheral blood mononuclear cells (PBMC) were isolated from single donor leukocyte cones purchased from the National Blood Transfusion Service (Colindale, UK) by density gradient centrifugation using histopaque gradient separation (Sigma-Aldrich, Catalog #: H8889). The obtained peripheral blood mononuclear cells were resuspended in RPMI-1640 medium, and monocytes were obtained by adhesion purification, after 2 h at 37°C and 5% CO_2_. The monolayer of the adherent peripheral blood mononuclear cells was washed 2x with HBSS to remove non-adherent cells. The cells were subsequently differentiated in human monocyte-derived macrophages (hMDMs) using 100 ng/ml macrophage colony-stimulating factor (PeproTech, catalog #:315-02) supplemented with 10% FBS, for 5 days. The macrophage purity was confirmed by immunohistochemical assessment of CD68.

### Cell stimulations

BMDMs were seeded in 24-well or 12-well plates at densities of 0.5×10^6^ and 1×10^6^ cells per well, respectively. THP-1 cells were seeded in 24-well or 12-well plates at densities of 1×10^6^ and 2×10^6^ cells per well, respectively. Following macrophage differentiation with PMA, THP-1 macrophages were washed and maintained in fresh PMA-free medium for an additional 24 h prior to any treatment. The concentrations of the stimuli used were selected based on prior studies demonstrating their ability to induce TLR signaling. For certain experiments (Figures 3, 4A, and 4B), BMDMs and THP-1 macrophages were treated overnight with either recombinant mouse IFNG (50 ng/ml; Sigma-Aldrich) or human IFNG (50 ng/ml; Sigma-Aldrich), respectively [29,30]. BMDMs and THP-1 macrophages were treated with LPS (500 ng/ml; Invivogen) for 6 h. THP-1 macrophages were stimulated with various TLR ligands; PAM3 (500 ng/ml; Invivogen), Poly I:C (1 μg/ml; Invivogen), and imiquimod (3 μg/ml; Invivogen) [31–34].

### Real-time quantitative PCR

Cells were lysed and RNA was isolated using TRIzol (T9424; Sigma-Aldrich) according to the manufacturer’s instructions. A total of 500 ng of RNA was then reverse transcribed in a Bio-Rad thermocycler using the High-Capacity cDNA Reverse Transcription Kit (4368814; Applied Biosystems), according to the manufacturer’s instructions. Real-time PCR was then performed using gene-specific primers **(Table 1)** and PowerUp SYBR Green Master Mix (A25741; Applied Biosystems). Real-time PCR was performed in an ViiA 7 Real-Time PCR System (Applied Biosystems).

**Table 1:**
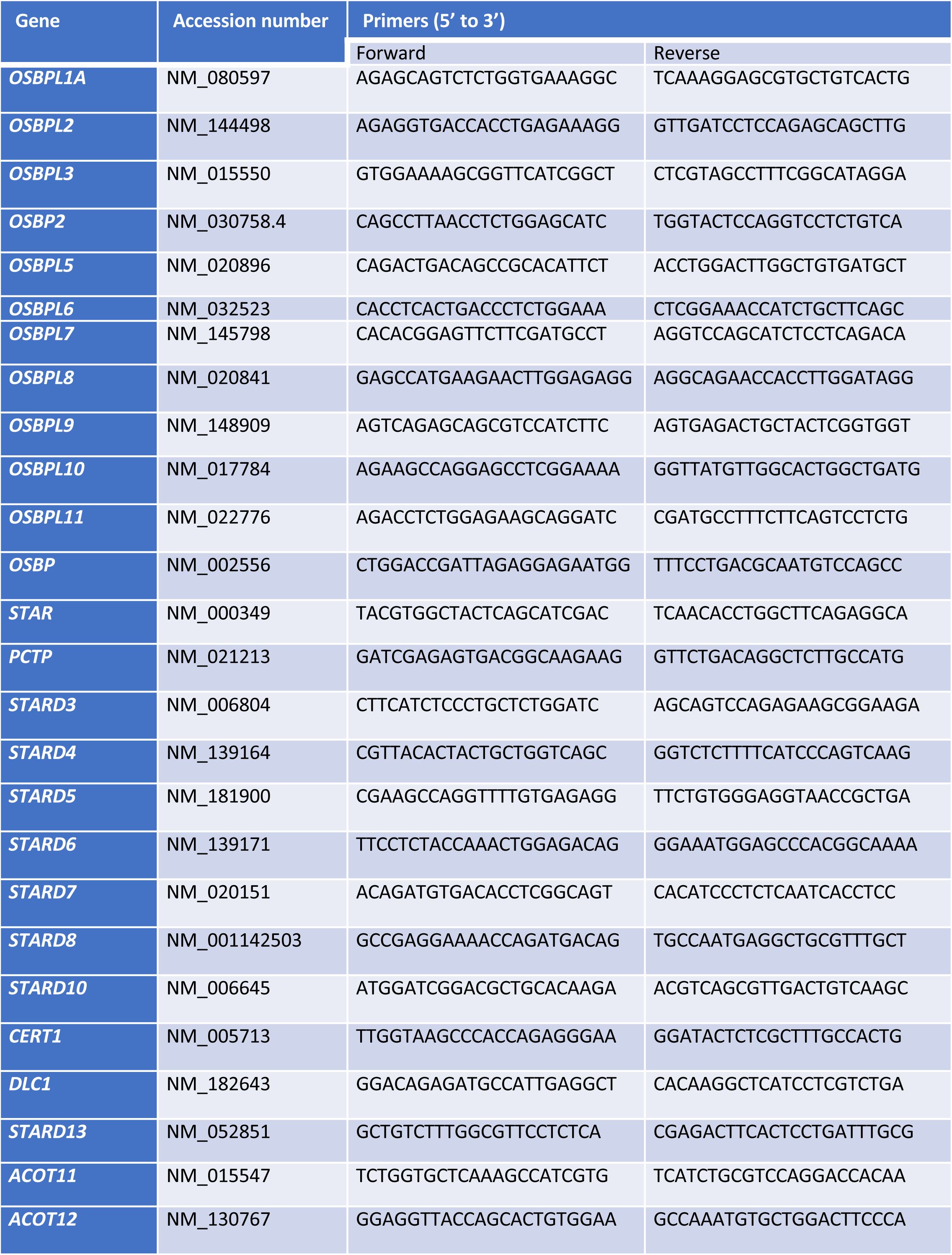
Primers used for RT-qPCR.

### RT-qPCR data analysis

The ΔΔCt algorithm [35] was used in order to relatively quantify the data obtained from the RT-qPCR experiments. GAPDH was used as a housekeeping gene for all experiments, as it was uniformly and constantly expressed in all samples, across all treatments. When comparing gene expression between hMDMs and THP-1 macrophages in Figure 1, no normalization methods were employed. This approach allowed us to directly compare gene expression differences while avoiding any potential biases introduced by data normalization methods. To maintain consistency in data acquisition, the PCR conditions were kept identical across species and throughout the study. The expression of all ORPs and STARDs were expressed in arbitrary units as stated in figure legends with or without log_10_ fold change (FC).

**Figure 1.**
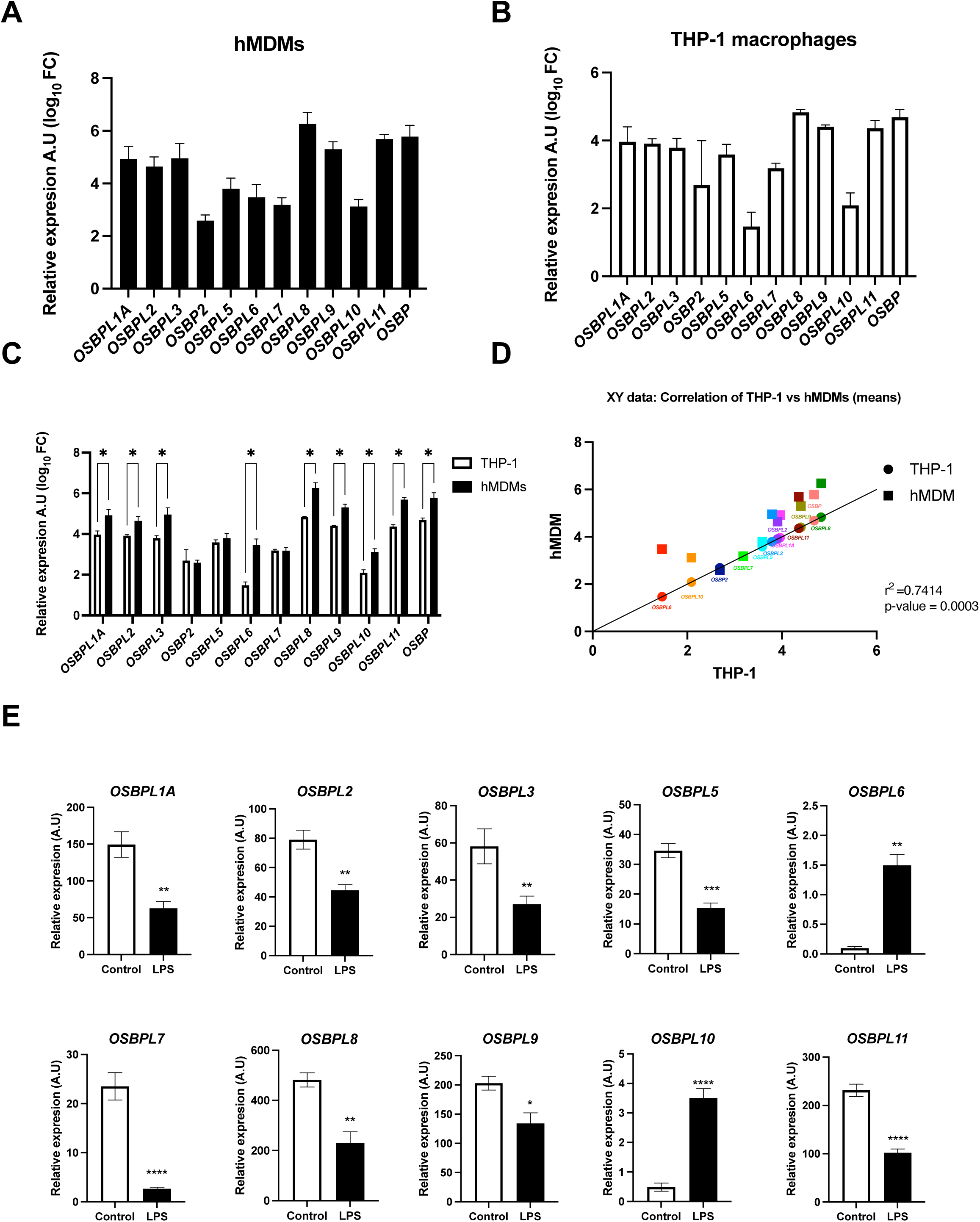
ORP proteins are constitutively expressed and are modulated in response to TLR ligation. Gene expression of all 12 ORP family members as determined by real-time qPCR in **(A)** hMDMs and **(B)** THP-1 macrophages. **(C)** Comparison of the expression of ORP family members between hMDMs and THP-1 -differentiated macrophages. **(D)** Correlation analysis of ORP family members between hMDMs and THP-1 -differentiated macrophages. **(E)** THP-1 differentiated macrophages were either left untreated or treated with LPS (500 ng/ml) for 6 h. RNA was extracted and converted to cDNA. Gene expression of all 12 ORP family members was determined by real-time qPCR and is shown as arbitrary units. Error bars represent SEM. The statistical analysis of the data was performed using the student’s t-test where *, *p*≤0.01; **, *p*≤0.001; ***, *p*≤0.0001; ****, *p*≤0.0001. In panel D, each ORP is denoted by a different color. In panel C, significance was calculated between each gene in the two cell types and in panel E, significance was calculated against the control, untreated group.

Statistics was performed using GraphPad Prism, by unpaired, two-tailed *t*-tests. The level of significance was set at *p*≤ 0.05 and where *, *p*≤0.01; **, *p*≤0.001; ***, *p*≤0.0001; ****, *p*≤0.0001. The number of replicates of each experiment is indicated in the figure legends (n=x). Values are given as mean ± standard error mean (SEM), from x independent experiments, as indicated.

### Data and code availability

All raw data reported in this paper will be shared by the lead contact upon request. This paper does not report original code.

Any additional information required to reanalyze the data reported in this paper is available from the lead contact upon request.

## Results

### ORP proteins are constitutively expressed and are modulated in response to TLR ligation

Lipids participate in various cellular functions and have been associated with inflammatory signalling. The distribution of lipids across cellular organelles is a key determinant of organelle function and signalling emanating from subcellular compartments. However, whether lipid transport proteins are regulated by immune signalling remains ambiguous. To investigate whether the individual members of the ORP family of proteins are constitutively expressed, we measured their expression in primary human monocyte-derived macrophages (hMDMs) by RT-qPCR. All ORP family showed detectable expression levels **(Fig. 1A)**. Correspondingly, THP-1 differentiated macrophages showed a similar expression pattern of ORP members, compared to that observed in hMDMs **(Fig. 1B)**. However, when comparing the two cell types, hMDMs displayed significantly higher expression levels for most ORPs, except *OSBP2, OSBPL5*, and *OSBPL7*, which showed comparable expression between hMDMs and THP-1 macrophages **(Fig. 1C)**. A correlation analysis confirmed a strong positive relationship in the expression of ORP family members between the two cell types (r² = 0.7414, p = 0.0003), validating THP-1 macrophages as a suitable model for subsequent experiments **(Fig. 1D)**.

Next, we investigated whether TLR4 signaling regulates the expression of ORP family members. THP-1 macrophages treated with LPS, a TLR4 agonist, for 6 hours exhibited significant downregulation of most ORP family members compared to untreated controls **(Fig. 1E).** Notably, the expression of *OSBP2* and *OSBP* was not altered (not shown) while the relative expression of *OSBPL6* and *OSBPL10* was significantly upregulated **(Fig. 1E)**. These results indicate that TLR4 signaling selectively and differentially regulates the expression of ORP family members.

### ORP expression is differentially regulated by TLR stimuli

Humans express 10 TLR family members which are involved in sensing a wide variety of pathogen-associated molecular patterns. These TLRs are predominantly localized at the plasma membrane with the exception of the TLRs 3, 7, 8 and 9 which are localized at the endosomal membrane. Distinct TLRs activate different signaling pathways via specific adaptor proteins, leading to unique gene expression profiles. For example, TLR4 predominantly signals through the MyD88 adaptor protein to activate NFKB and MAP-kinase pathways but can also signal via TRIF, under certain conditions [36–39]. In contrast, TLR3 exclusively signals through TRIF to induce the expression of type-I interferons [40].

To determine whether the observed regulation of ORPs is specific to TLR4 or also initiated by other TLRs, THP-1 macrophages were exposed to ligands for TLR2, TLR3, and TLR7, in addition to TLR4. Consistent with our previous results (**Fig. 1E**), exposure to LPS resulted in a significant downregulation of the majority of the ORP family members, with the exception of *OSBP2* and *OSBP* **(Fig. 2)**. By contrast, *OSBPL6* and *OSBPL10* exhibited an increase in expression. Stimulation with the TLR2 ligand, Pam3CSK4, significantly downregulated *OSBPL1A, OSBPL7, OSBPL9, OSBPL10, OSBPL11,* and *OSBP* while other ORPs were unaffected. Poly(I:C), a TLR3 ligand, suppressed the expression of *OSBPL1A*, *OSBPL2*, *OSBPL5*, *OSBPL7*, *OSBPL8*, *OSBPL9*, *OSBP11*, and *OSBP*, while imiquimod, a TLR7 ligand, had no significant effects on any of the ORPs **(Fig. 2)**. This data summarizing changes in ORP gene expression by TLRs is presented in **Table 2**. These results suggest that the regulation of most ORPs is independent of TRIF-mediated signaling. Moreover, the specific regulation of most ORPs by TLR4, but not TLR2, suggests the involvement of additional factors beyond the shared downstream pathways between the two receptors.

**Figure 2.**
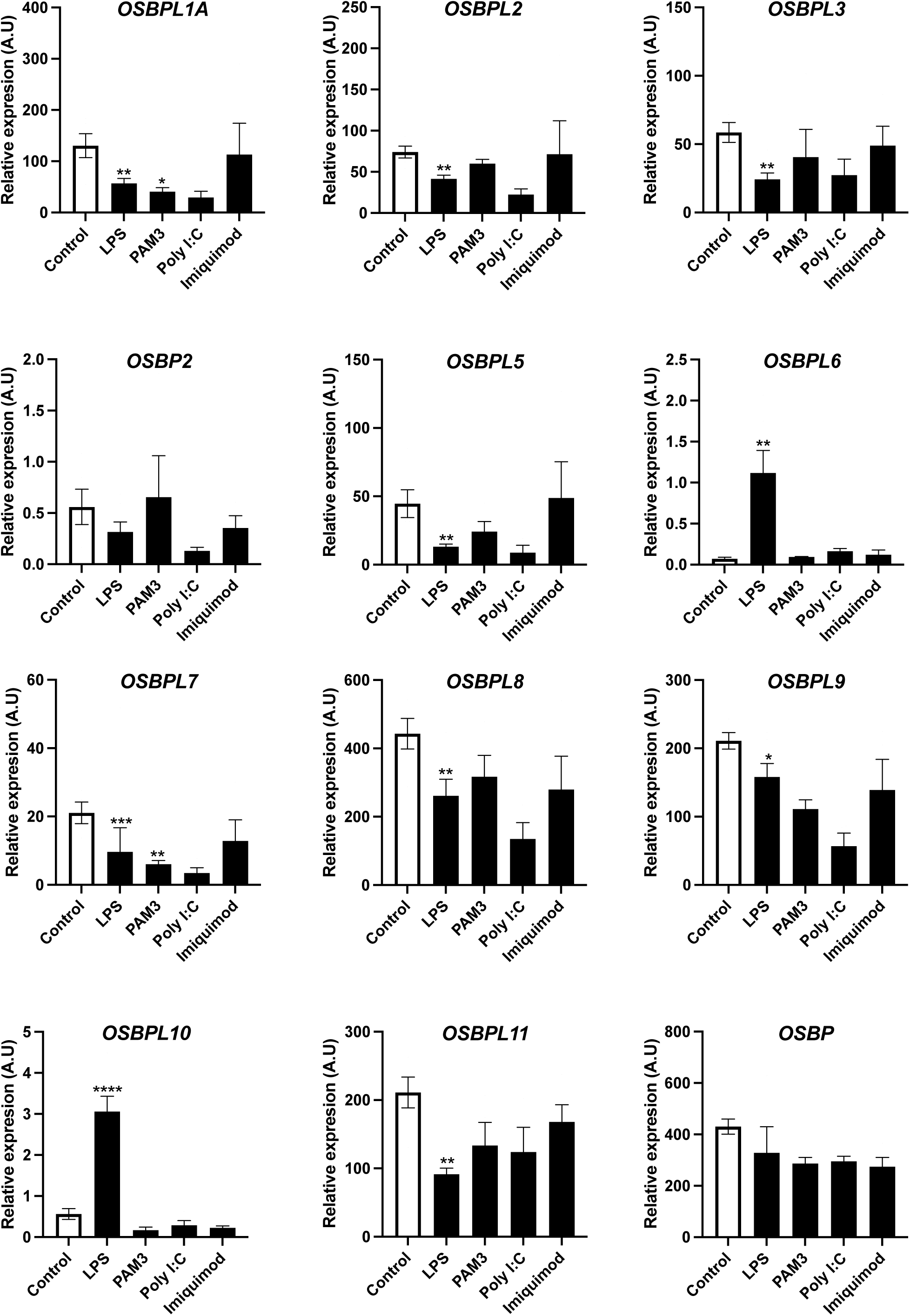
ORP expression is differentially regulated by TLR stimuli. THP-1 differentiated macrophages were either left untreated or treated with LPS (500 ng/ml), PAM3 (500 ng/ml), poly I:C (1 μg/ml), or imiquimod (3 μg/ml) for 6 h. RNA was extracted and converted to cDNA. Gene expression of all 12 ORP family members was determined by real-time qPCR. The gene expression of ORP family members upon various treatments is shown as arbitrary units, n=4. Error bars represent SEM. The statistical analysis of the data was performed using the student’s t-test where *, *p*≤0.01; **, *p*≤0.001; ***, *p*≤0.0001; ****, *p*≤0.0001. Significance was calculated against the control, untreated group.

**Table 2.**
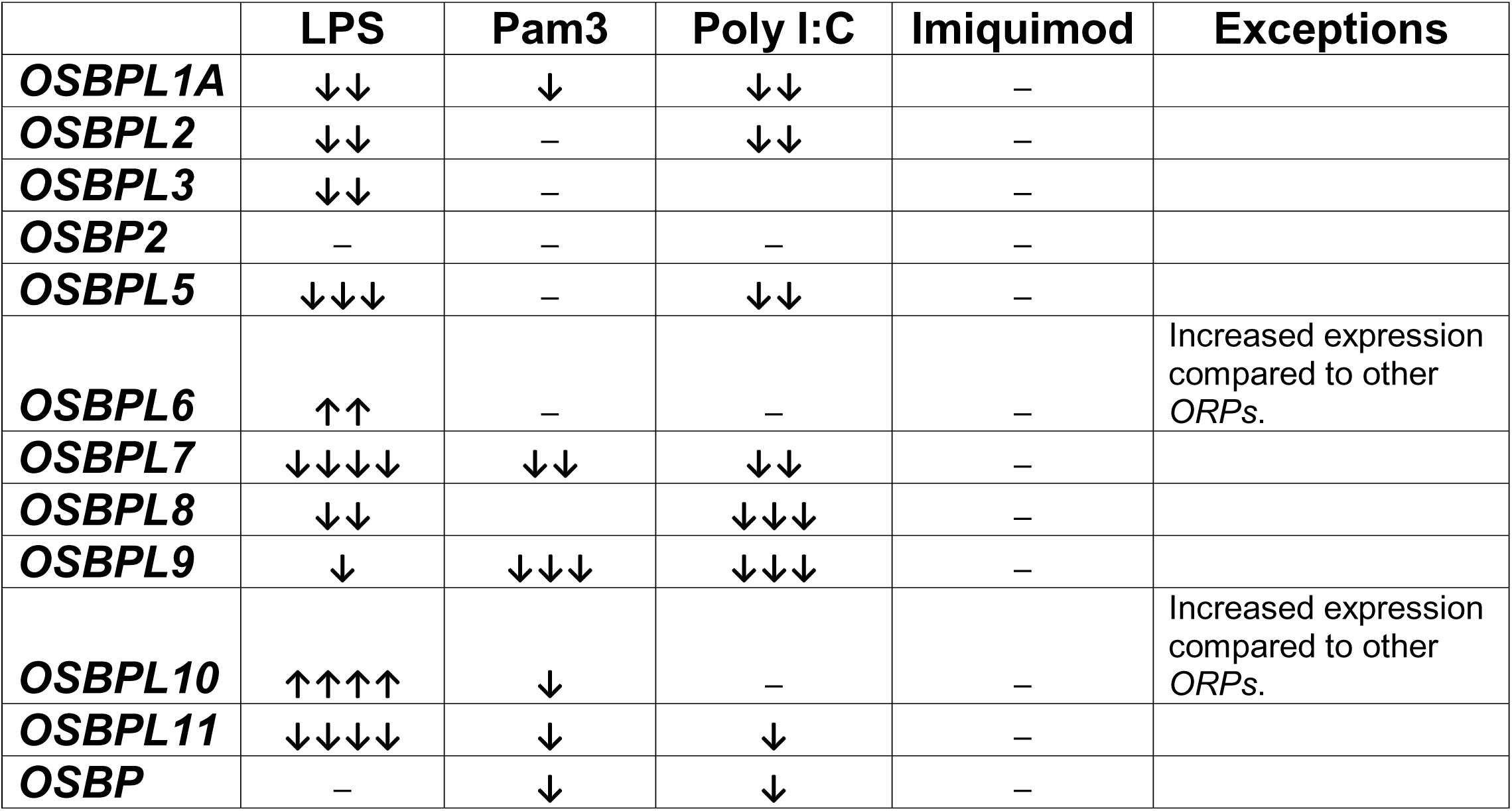
Changes in ORP expression in response to different TLR ligands in THP-1 macrophages. Arrows indicate direction of change. Arrows pointing upwards show an increase while downward facing arrows show a decrease in gene expression. The number of arrows correspond to the level of significance (e.g., ** for two arrows). A dash indicates no significant change in gene expression.

### TLR signaling does not regulate the expression of STARD family members

Lipid transport is essential to lipid metabolism facilitating the movement of lipids between cellular compartments and enabling energy storage and cellular signaling. Beyond ORPs, STARD proteins also play key roles in intracellular lipid transport. Consisting of 15 family members, STARD proteins transport cholesterol and phospholipids across organelles, which is important to mitochondrial function and cellular homeostasis. To investigate whether TLR signaling regulates STARD proteins, we analyzed their expression in THP-1 macrophages. Unlike ORPs, STARD gene expression remained unaltered, with the exception of *STARD10*, following TLR4 activation with LPS (**Fig. 3**). Although *STARD9* could not be assessed due to inefficient primers, the lack of regulation in other STARD family members suggests that TLR4 signaling specifically targets ORPs without affecting STARD proteins.

**Figure 3.**
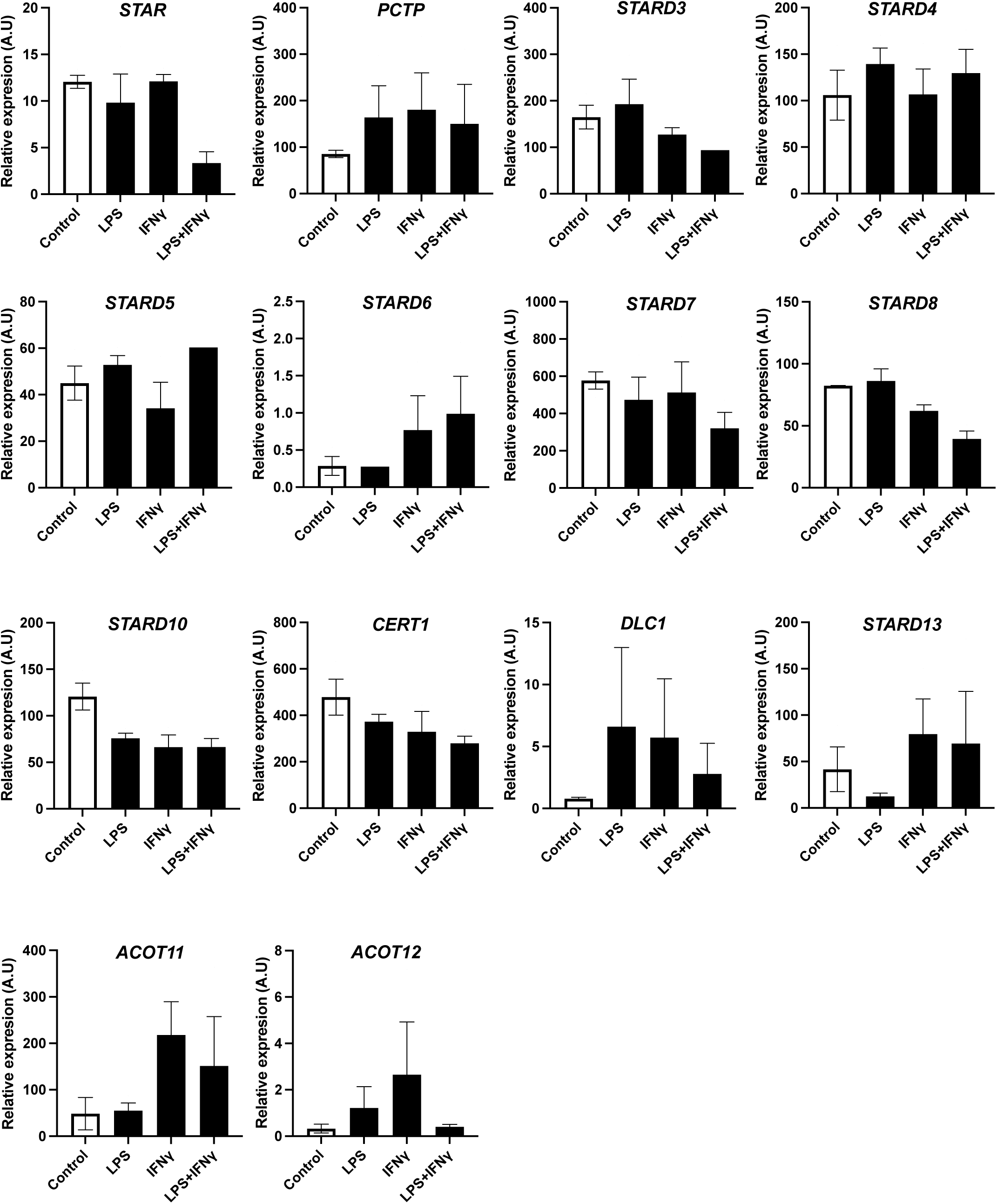
TLR signaling does not regulate the expression of STARD family members. THP-1 differentiated macrophages were either left untreated or treated with LPS (500 ng/ml) for 6 h, human IFNG (50 ng/ml) overnight, or a combination of both. RNA was extracted and converted to cDNA. Gene expression of 14 STARD family members was determined by real-time qPCR. The expression of the STARD family members upon various treatments is shown as arbitrary units, n=4. Error bars represent SEM. The statistical analysis of the data was performed using the student’s t-test. Significance was calculated against the control, untreated group.

IFNG is a crucial innate immune stimulus, which has previously been shown to modulate lipid metabolism and transport. Specifically, IFNG inhibits genes involved in cholesterol and fatty acid synthesis pathways. Moreover, when combined with LPS, IFNG has the ability to enhance lipid efflux and promote lipid catabolism [5]. Considering these roles, we next examined whether IFN-γ by itself or in combination with LPS has the ability to regulate *STARD* genes. To investigate this, THP-1 macrophages were exposed to IFNG overnight in the absence or presence of prior LPS stimulation for 6 hours. After overnight stimulation, purified RNA was subjected to RT-PCR. Among the STARD family members, STARD1, STARD8, and STARD10 exhibited modest downregulation in response to the combined IFNG and LPS treatment, while other members showed no significant changes (**Fig. 3**). These findings suggest that STARD genes are not as prominently regulated by immune stimuli as the ORP family members, indicating a distinct regulatory framework for these two protein families.

### Species-specific differences in ORP regulation by LPS and IFNG

Our results so far indicate that ORP genes are selectively regulated by TLR signaling, while STARD genes are not affected by either TLR or IFNG signaling. To further investigate whether IFN-γ modulates ORP expression, we exposed human THP-1 macrophages to IFNG alone or in combination with LPS. Consistent with our previous findings, LPS alone resulted in significant downregulation of most ORPs. However, IFNG alone had no significant effect on ORP expression, and its combination with LPS produced a similar expression profile to LPS alone. These results suggest that IFNG does not play a role in modulating the expression of lipid transport genes in human macrophages.

Previous studies have highlighted specific differences in lipid transport and metabolism between humans and mice. For example, the expression of cholesteryl ester transfer protein, an enzyme that facilitates cholesterol transfer between lipoproteins, significantly differs between mice and humans [41,42]. Similarly, the gene expression of *APOA1*, which is associated with HDL metabolism, differs between the species[43]. To explore species-specific differences, we next examined ORP expression in mouse bone marrow-derived macrophages (BMDMs) under the same conditions (**Fig. 4**). Unlike in human cells, IFNG alone significantly downregulated *Osbpl2*, *Osbpl8*, *Osbpl10*, and *Osbp* in mouse BMDMs (**Fig. 4A, B**). LPS treatment in BMDMs also regulated a more limited subset of ORPs, specifically downregulating *Osbpl1*, *Osbpl2*, and *Osbpl10*. Interestingly, human *OSBPL10* exhibited an increase in expression following LPS treatment **(Fig. 2)**, underscoring distinct regulatory mechanisms between species. When IFNG and LPS were combined in BMDMs, only a limited synergistic effect was observed, particularly for *Osbpl1* and *Osbpl10*. A descriptive heatmap summarizes these species-specific differences, highlighting the contrasting effects of LPS and IFNG on ORP expression in human THP-1 macrophages and mouse BMDMs (**Fig. 4C**). These findings emphasize the importance of species context when investigating immune-regulated lipid metabolism and transport.

**Figure 4.**
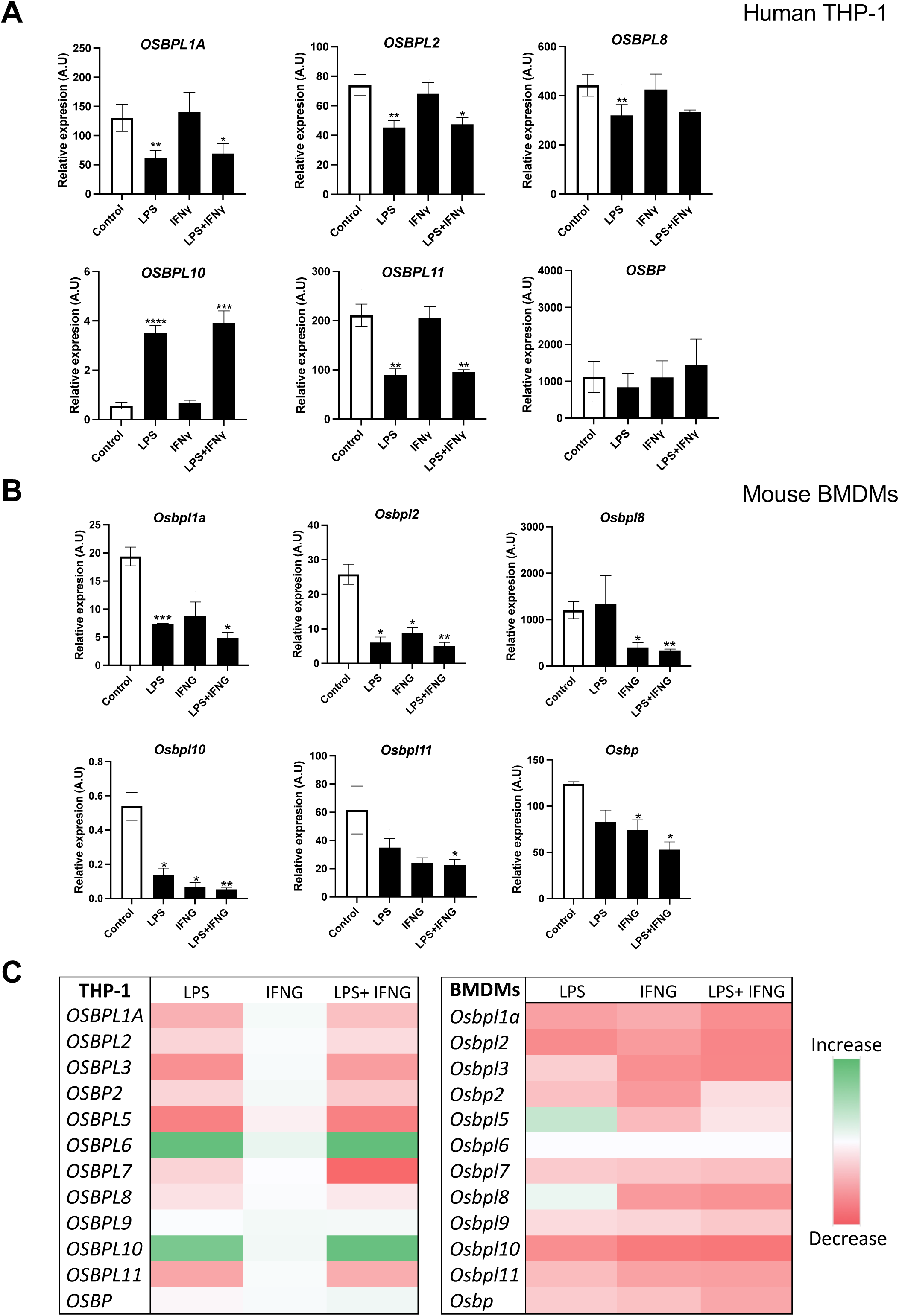
Species-specific differences in ORP regulation by LPS and IFNG. BMDMs and THP-1 differentiated macrophages were either left untreated or treated with LPS (500 ng/ml) for 6, human or mouse IFNG (50 ng/ml) overnight, or a combination of both. RNA was extracted and converted to cDNA. Gene expression of all 12 ORP family members was determined by real-time qPCR. The relative expression of only the significant ORPs differentially regulated by IFNG in **(A)** BMDMs was plotted alongside the expression of these ORPs in **(B)** THP-1 differentiated macrophages. Relative expression of ORPs is shown as arbitrary units. **(C)** Descriptive statistics were performed on the effect of each treatment against the control, for each ORP member in BMDMs, and the results were plotted in a heat map. Red shading indicates a decrease, while green shading represents an increase. A darker shade of red or green color signifies a greater decrease or increase, respectively. n=8 for THP-1 cells and n=4 for BMDMs. Error bars represent SEM. The statistical analysis of the data was performed using the student’s t-test where *, *p*≤0.01; **, *p*≤0.001; ***, *p*≤0.0001; ****, *p*≤0.0001. Significance was calculated against the control, untreated group.

## Discussion

Lipid transport proteins, such as the ORPs and STARDs families, are central to cellular lipid metabolism and contribute to lipid transport across organelles. In this study, we report that the lipid transporters are regulated by immune stimuli in human and mouse macrophage models. Our results underscore the complexity and specificity of immune regulation on lipid transporters with distinct patterns observed between the ORP and STARD family members.

Our findings demonstrate that all 12 members of the ORP family are constitutively expressed and depict a positive correlation between hMDMs and THP-1 macrophages. However, their expression levels vary between the two cell types with generally higher expression levels observed in hMDMs. TLR4 activation through LPS resulted in significant downregulation of most ORP members in THP-1 macrophages, except for *OSBPL6* and *OSBPL10*, which were upregulated. Notably, stimulation of TLR2, TLR3, and TLR7 also modulated specific ORPs, albeit to a lesser extent. For example, Pam3CSK4 (TLR2 ligand) reduced *OSBPL1A*, *OSBPL7, OSBPL9,* and *OSBPL11* expression, while poly(I:C) (TLR3 ligand) and imiquimod (TLR7 ligand) primarily affected *OSBPL7* and *OSBPL11*. These data suggest that TLR4 has a broader impact on ORP regulation compared to other TLRs but highlight the nuanced and context-dependent nature of ORP modulation by different immune pathways. These results are consistent with published findings showing that LPS alters the expression of genes involved in lipid metabolism [13,15]. The upregulation of *OSBPL6* and *OSBPL10* suggests that these proteins may have compensatory or specialized roles in maintaining lipid homeostasis during immune activation. Future work could focus on elucidating their specific lipid substrates or roles in immune-metabolic interactions.

Despite the established role of TLRs in regulating lipid metabolism, the specificity of interactions between different TLRs and lipid transport proteins such as ORPs and STARDs remains largely unexplored. While our findings reveal distinct regulatory patterns for ORPs across TLR2, TLR3, TLR4, and TLR7 signalling, the molecular basis of these specificities remains unknown. Future studies are needed to investigate how different TLRs uniquely influence lipid transport proteins and how these pathways contribute to immune and metabolic homeostasis.

Interestingly, these responses were specific to TLR4 ligation, as none of the other TLR ligands tested such broadly altered ORP expression. This suggests that TLR4-mediated ORP regulation is not solely due to shared downstream pathways but may involve additional factors unique to TLR4 signalling.

Innate immune agonists activate distinct but overlapping transcriptional programs that provide a plausible framework for the stimulus-specific regulation we observe across lipid-transport gene families. Downstream of TLR ligands (including LPS/TLR4), MyD88- and TRIF-linked signalling rapidly engages canonical inflammatory transcription factors such as NFKB and AP-1, with additional involvement of IRF-family factors depending on the cellular context [47,48]. In parallel, IFNG signals through the JAK–STAT pathway to activate STAT1 and induce IRF1, and many IFNG–responsive genes rely on combinatorial control (e.g., STAT1/IRF1 cooperation) encoded by promoter and enhancer cis-elements [49]. Notably, TLR4 signalling has been reported to engage IRF1 in macrophages and to promote the activation of IFN-stimulated loci, offering a mechanistic precedent for LPS-driven regulation of gene subsets that overlap with IFN-like transcriptional programs [50]. Consistent with these observations, the endosomal TLR ligands tested in this study did not reproduce the broad ORP repression observed with LPS. Since TLR7 signals via MyD88, this suggests that MyD88 signalling alone is not sufficient to drive widespread ORP transcriptional changes in this context and instead points to TLR4-specific features (e.g., receptor compartmentalization and/or additional downstream signalling inputs) as contributors. These immune transcription factor pathways can also directly intersect with lipid metabolism and reshape lipid-handling gene expression. For example, IFNG can downregulate LXR–ABCA1–linked cholesterol efflux programs in macrophage models via a JAK/STAT1-dependent mechanism [51]. Together, these precedents support a model in which differential responsiveness of ORP vs STARD genes reflects differences in immune-responsive cis-regulatory features and/or chromatin accessibility at these loci. The species-specific effects could potentially arise from divergence in transcription factor usage or regulatory elements. These possibilities can be tested in future work by targeted transcription factor occupancy assays or focused reporter approaches.

In contrast to the observed regulation of the ORP family, the STARD family, including well-studied lipid transporters like *STAR, PCTP*, and *STARD3*, were less responsive to immune stimuli. While most STARD members were unaffected by LPS or IFNG stimulation, combined treatment with LPS and IFNG led to modest suppression of *STAR*, *STARD8*, and *STARD10* expression. This limited response contrasts with the broad regulation observed in the ORP family. A plausible hypothesis is that the STARD proteins are regulated by metabolic or homeostatic pathways rather than immune signalling, consistent with their basal roles in organelle-specific lipid shuttling [52]. These findings indicate a potential functional divergence between the ORP family members and the STARD family members, with the former more actively engaged in immune-related lipid transport and the latter more aligned with basal metabolic cellular processes.

When comparing human and mouse macrophages, we observed notable species-specific differences in ORP expression. Mouse BMDMs displayed a more limited response to LPS compared to human THP-1 macrophages, with only *Osbpl1a*, *Osbpl2*, and *Osbpl10* being downregulated. Conversely, human *OSBPL10* expression was increased following LPS stimulation, suggesting species-specific differences in regulatory mechanisms. Additionally, IFNG alone downregulated *Osbpl2*, *Osbpl8*, *Osbpl10*, and *Osbp* in mouse BMDMs, a response that was not observed in human cells. These results are consistent with studies demonstrating variations in lipid metabolism between human and murine models [43,53], and may reflect evolutionary adaptations in lipid metabolism and immune function. These differences underscore the complexity of species-specific regulation, complicate the extrapolation of murine data to human immunological studies, and emphasize the importance of validating findings from murine models in human systems. Further comparative studies across species are essential for identifying both conserved and species-specific regulatory mechanisms, providing critical insights that can guide translational applications in immunology and pharmacology. However, comparative studies pose challenges, such as differences in baseline gene expression, species-specific immune adaptations, and variations in experimental protocols, which must be carefully addressed.

The regulation of lipid transport by immune signalling is critically relevant to several disease contexts, yet it remains inadequately studied. Dysregulated lipid metabolism contributes to the pathogenesis of diseases such as atherosclerosis, diabetes, and chronic inflammatory disorders, where the interaction between immune signalling and lipid transport mechanisms is poorly understood. Our findings suggest that ORPs, which are selectively regulated by TLR4 signalling, could represent novel therapeutic targets for such conditions. Integrating molecular mechanisms into disease contexts, particularly in metabolic and cardiovascular diseases, will enhance the translational impact of future research.

In conclusion, this study provides novel insights into the selective regulation of lipid transport proteins by immune signalling. The differential regulation of the ORP family members by TLR4 and the lack of modulation of the STARD family members suggest distinct roles for these protein families in immune and metabolic contexts. Furthermore, the observed species-specific differences in ORP regulation underscore the importance of considering species-context when interpreting immune-metabolic interactions. Future work should focus on elucidating the molecular mechanisms underlying these regulatory differences, including defining the transcription factor programs and cis-regulatory features that drive the stimulus-and species-specific ORP and STARD regulation observed in this study.

