## Supplementary figures and images for "Distinct lipid transport proteins are regulated by innate immune stimuli"

### Fig. S1

#### Figure S1. Domain architecture and classification of ORPs

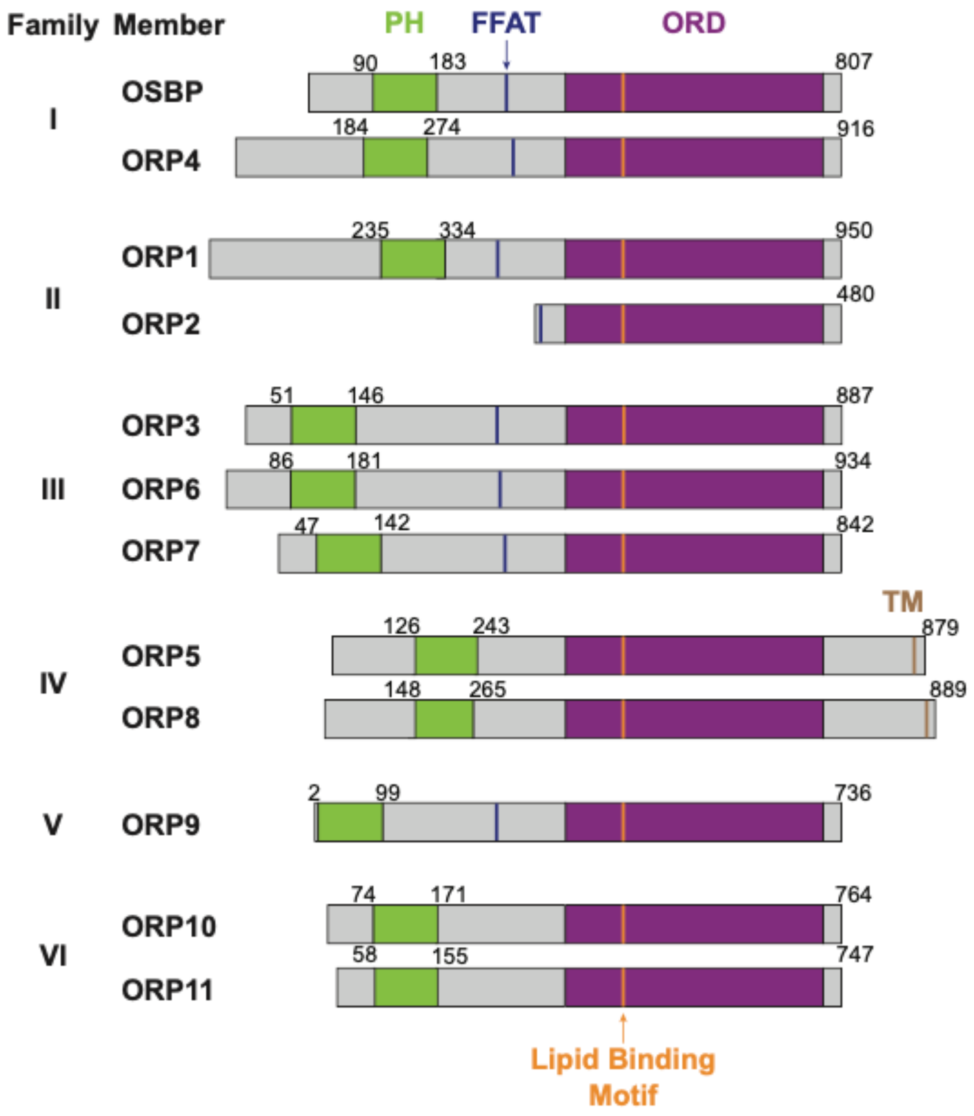


**Figure S1**
